# Depression reduces structurally informed network flexibility in premanifest Huntington’s disease

**DOI:** 10.1101/2025.09.25.678149

**Authors:** Tamrin Barta, GradDipAdvPsych, Matthew D. Greaves, GradDipAdvPsych, Leonardo Novelli, Yifat Glikmann-Johnston, Adeel Razi

**Affiliations:** Turner Institute of Brain and Mental Health at the School of Psychological Sciences, and, Faculty of Medicine, Nursing and Health Sciences, Monash University; Monash Biomedical Imaging, Monash University, Clayton, Victoria, Australia; Queen Square Institute of Neurology, University College London, London, United Kingdom; CIFAR Azrieli Global Scholars Program, CIFAR, Toronto, Ontario, Canada

## Abstract

1

**Background and objectives:** The extent to which structural connectivity constrains effective connectivity in both depression and neurodegenerative contexts remains poorly understood. In particular, the relationship between structural connectivity aberrations and effective dysconnectivity associated with depression in Huntingtin’s disease remains uncharacterized. Here, we applied a novel procedure that implements structural connectivity-informed spectral dynamic causal modelling to examine how structural connectivity shapes directed inter-regional influences in premanifest Huntington’s disease gene expansion carriers (HDGECs) with and without depression history.

**Methods:** Using spectral dynamic causal modeling embedded in a hierarchical empirical Bayes framework, we analyzed fMRI data from 98 premanifest HDGECs across default mode network and striatum (caudate and putamen). HDGECs were split into two groups based on either having a history of depression or not. Depression severity on both the Beck Depression Inventory, 2nd Edition (BDI-II) and Hospital Anxiety and Depression Scale, Depression Subscale (HADS-D) was used to measure clinically elevated depression symptoms. Leave-one-out cross-validation was implemented to test predictive validity.

**Results:** Model evidence substantially favored structurally informed over uninformed approaches across all participants. For HDGECs, having a history of depression was associated with reduced baseline variability in effective connectivity (decreased α parameter), with particularly tight regularization of near-zero-valued structural connections toward zero effective connectivity values while leaving strongly connected pathways relatively unaffected. Effects converged on striatal self-connectivity and hippocampal-striatal pathways, with distinct patterns emerging between depression history groups. Notably, clinically elevated depression revealed differential connectivity signatures, with right caudate self-connectivity showing positive correlations with clinical cut-offs for HDGECs with and without depression history. In leave-one-out cross-validation, specific connections including DMN-to-striatum (BDI: *r* = -0.31, *p* = .002; HADS-D: *r* = -0.33, *p* = .001), right hippocampus-to-left caudate (BDI: *r* = -0.46, *p* < .001; HADS-D: *r* = -0.30, *p* = .002), and left caudate-to-left putamen (BDI: *r* = -0.48, *p* < .001; HADS-D: *r* = -0.30, *p* = .003) significantly predicted individual differences in depression severity scores.

**Discussion:** Together, these findings link reduced network flexibility to depression vulnerability in premanifest neurodegeneration, providing a mechanistic bridge between anatomical constraints, effective connectivity alterations, and clinical depression phenotypes.

## 2 Introduction

Huntington’s Disease (HD) is a monogenetic neurodegenerative disease that is unique in its autosomal dominant inheritance(1) and protracted course, where people typically develop progressive motor, cognitive, and psychiatric symptoms in mid-life(2,3). Depression represents a predominant psychiatric manifestation in HD, with meta-analytic pooled prevalence rates of 23% in premanifest HD gene expansion carriers (HDGECs; before clinical diagnosis based on motor signs) and 38% in manifest (clinically diagnosed) HD(4), equating to 2–3.5–fold elevation when compared to the general population(5). Prevalence varies across studies but is consistently elevated, with point prevalence ranging from 15–52% and lifetime prevalence of 20% in premanifest HDGECs(6,7). Depressive symptoms peak just prior to manifest diagnosis and in early manifest stages(8). In the brain, early and severe degeneration of the striatum is pathognomonic, occurring from the premanifest period(9–12). Outside the striatum, cortical volume changes emerge during the transition to manifest diagnosis(11), with cortical atrophy becoming most pronounced 20 years after manifest onset(13). Despite characterization of the neurodegenerative process and depression trajectories throughout HD progression, structural atrophy severity does not explain variation in depression symptomatology or severity. Specifically, whole-brain and regional volume loss demonstrate no association with affective changes(14), and basal ganglia atrophy does not correlate with depressive symptoms(15). Depression in HD has been more consistently associated with dysconnectivity within mood-regulatory networks, including cortico–striatal pathways and default mode network (DMN)(16–18).

A fundamental question arises how depression and HD pathogenesis disrupt or influence the structure– function relationships. In healthy brains, structural and functional connectivity demonstrates significant but loose coupling, with anatomy constraining but not determining functional dynamics(19). Studying the quantitative relationship between structural and functional connectivity—often termed structure–function coupling—has emerged as a means to understand disease-related network reorganization(20). Structure– function coupling quantifies the degree of correspondence between structural and functional network organization, providing a framework for understanding how HD pathogenesis disrupts normal brain network relationships through mechanisms such as neural network compensation, where functional connectivity is maintained despite structural degeneration(21,22), and pathological hyperconnectivity, where functional activity increases independent of (or contrary to) underlying structural connectivity loss(23). The fluctuating temporal pattern of depression in HD may be explained by changes in structure–function dynamics over HD progression. For example, coupling may decrease due to neurodegeneration and white matter injury in surrounding fiber tracts(24). Traditional approaches to investigate structure–function coupling rely on post-hoc correlational analyses between independently estimated structural and functional connectivity measures. Recent methodological advances integrate structural connectivity information directly into the modeling procedure for effective connectivity estimation, providing a mechanistic framework for understanding how structural constraints systematically influence functional network dynamics(25). In HD, significant white matter dysconnection in interhemispheric connections in frontal and parietal cortices is shown in manifest HD(26), and stronger structural connectivity has been associated with weaker functional activity—and vice versa—alongside antero–posterior dissociation of functional connectivity in premanifest HDGECs(27). There is evidence that depression in premanifest HDGECs is associated with increased functional connectivity within default mode network (DMN) regions (but not striatum) and decreased structural connectivity between DMN and basal ganglia(17). Despite these emerging findings, the mechanisms governing structure–function relationships remain incompletely characterized, as does the causal interactions reflected in effective connectivity measures(25), particularly regarding how the neurodegenerative process systematically modifies these network properties.

Structure–function dynamics are altered in other neurodegenerative diseases and in major depression, but the pattern of alterations remains inconsistent across studies. There is evidence of progressive structure– function decoupling (weakening correspondence) in Alzheimer’s disease(28) and Parkinson’s disease(29), as well as conflicting evidence of increased structure–function coupling in DMN in Alzheimer’s disease(30)—suggested to represent functional collapse to structure, whereby blood-oxygen-dependent (BOLD) correlations increasingly track the anatomical connectome, indicating reduced network flexibility and a narrower dynamical repertoire (i.e., less integration, fewer state transitions, more reliance on monosynaptic topology). In major depression, structure–function decoupling has been positively associated with depression severity(31) but, similar to Alzheimer’s disease, increased structure–function coupling has been shown in the DMN associated with major depression(32). Taken together, changes in structure– function organization appear to be network specific, with increased coupling of DMN associated with both neurodegeneration and depression. Given the focus on functional reorganization as an explanatory mechanism in HD(21,22), a critical open question is when (and in whom) structure–function dynamics undergo fundamental reorganization.

Here, we leverage the largest dataset of premanifest HDGECs including diffusion and functional MRI, TrackOn-HD(14,22), to apply non-generic structural connectivity priors to inform effective connectivity analyses. Although existing methods examine correlations between nodal connectivity profiles(28,31) or employ harmonic decomposition to assess structural constraints on functional dynamics(33,34), these approaches cannot capture the directional causal interactions (i.e., effective connectivity) fundamental to network communication. Our framework addresses this limitation by using structural connectivity to systematically scale the precision of effective connectivity estimates. To overcome computational infeasibility of optimizing priors for each participant(35), we apply a novel hierarchical empirical Bayes (HEB) procedure(36) where group-level structural constraints inform causal interactions in spectral dynamic causal models (spDCM)(37–39). This methodological integration represents a novel form of structure– function coupling that quantifies how white matter architecture directly influences the reliability and directionality of neural communication patterns of resting-state dynamics.

We examine how structurally informed effective connectivity patterns differ across regions and investigate whether HD pathogenesis fundamentally reorganizes these relationships associated with depressive symptomatology at the group level. This approach represents a novel application of structure–function coupling principles, where structural connectivity precision parameters (the degree to which structural connection strength allows the model to shrink effective connectivity estimates toward anatomical connections) systematically modulate effective connectivity estimates across participant groups, extending beyond traditional correlational methods to capture directional network communication under anatomical constraints. Drawing from convergent evidence, we hypothesize that HDGECs with a history of depression will demonstrate increased structural scaling of effective connectivity precision compared to HDGECs with no depression history, indicative of tighter constraint of functional dynamics around structural predictions. We anticipate that HDGECs with depression history will exhibit increased group level effective connectivity within striatal networks and altered DMN-striatal communication patterns, reflecting pathological reorganization.

## 3 Methods and Materials

### 3.1 Participants

The study included baseline clinical and imaging data from the existing TrackOn-HD study(22). TrackOn-HD was approved by local ethics committees at all participating sites and participants provided written informed consent in accordance with the Declaration of Helsinki. All participants had a cytosine, adenine, guanine (CAG) repeat length greater than 39. Participants were categorized into two groups based on history of depression (current and remitted) or no history of diagnosed depression. Exclusion criteria are reported in supplementary methods and depression history classifications are reported in supplementary results (SI.Table 2).

### 3.2 Clinical Measures

The study included two self-report measures of depressive symptoms: the Beck depression inventory, 2nd edition (BDI-II)(40) and the hospital anxiety and depression scale, depression subscale (HADS-D)(41). Clinical cut off scores developed for HD(42) distinguished clinically elevated depression symptom levels on the BDI-II (cut off of 10/11) and HADS-D (6/7). TrackOn-HD involved a comprehensive neuropsychiatric battery, including demographic and clinical measures. The study controlled concomitant regular (daily) mood medication use, including antipsychotics, benzodiazepines, selective serotonin reuptake inhibitors (SSRIs), and non-SSRIs. Medication type, regime, and indication are reported in supplementary results (SI.Table 3).

### 3.3 Statistical Analyses

See Fig. 1d–i for an outline of the modelling pipeline, described in detail below.

**Fig. 1.**
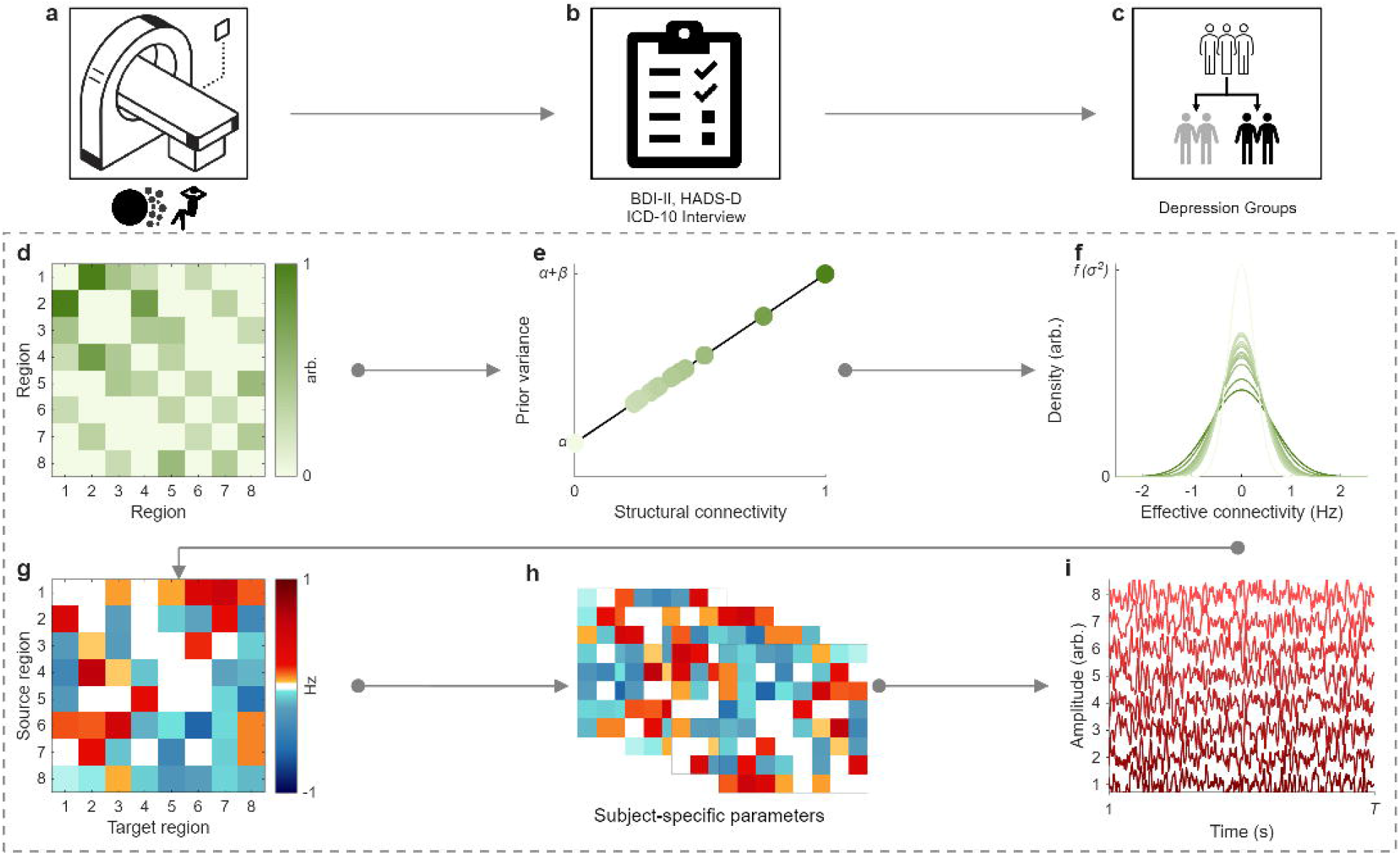
Procedure for evaluating structure–function relationships. **(a–c)** Study protocol, where **(a)** 3T resting-state and diffusion-weighted MRI are acquired, **(b)** key variables were ICD-10 categorization for depression history and Beck depression inventory, 2nd edition (BDI-II) and hospital anxiety and depression scale, depression subscale (HADS-D) symptom severity measurement, and **(c)** participant groups were based on depression history status. **(d–i)** In the hierarchical empirical Bayes model, normalized structural connectivity weights **(d)** are linearly transformed with intercept α and slope β to define prior variances over effective connectivity parameters **(e)**, such that greater structural connectivity corresponds to broader priors when β>0 (allowing for larger range of effective connectivity). These priors specify the distribution over effective connection strengths **(f)**, from which a group-level effective connectivity is drawn **(g)**. Subject-specific effective connectivity **(h)** captures individual deviations from this group-level model. At the individual level, the DCM forward model generates predicted blood-oxygen-dependent (BOLD) time series **(i)** from effective connectivity parameters.

#### 3.3.1 Data Acquisition

3T MRI data were acquired on two scanner systems at four sites: Philips Achieva (Vancouver and Leiden) and Siemens TIM Trio (London and Paris). Acquisition parameters are reported in supplementary methods.

#### 3.3.2 Regions of Interest

The selection of regions of interest (ROIs) and their size was based on previous research(39,43,44). ROIs and MNI coordinates were MPFC [3,54,-2], PCC [0,-52,26], hippocampus (left [-29,-18,-16], right [29,-18,-16]), caudate nucleus (left [-10,14,0], right [10,14,0]), and putamen (left [-28,2,0], right [-28,2,0]). Each ROI included an 8 mm sphere for MPFC and PCC and a 6 mm sphere for all other regions.

#### 3.3.3 MRI Pre-processing

DWI data was preprocessed using MRtrix3 (version 3.0.3)(45), FSL (version 6.0.4)(46) and ANTs (version 2.4.3)(47). Given the absence of reverse phase-encoding acquisitions, susceptibility distortions were corrected using Synb0-DisCo (v3.1)(48). Quality assurance was performed by two independent researchers. Fiber orientation distributions were reconstructed in participant native T1 space using the iFOD2 algorithm(49), and anatomically constrained tractography was used to dynamically seed and generate 4 million streamlines per subject(50,51). Streamlines were parcellated using a custom atlas comprising spherical ROIs (further information, supplementary methods).

Resting-state fMRI pre-processing was performed using *fMRIPrep* v21.02.2(52,53) and MRIQC v22.0.6(54), using FreeSurfer v-6.0.1(55), as previously reported(18). See Supplementary Material for full *fMRIPrep* pipeline. ROI time series was calculated as the first principal component of the voxels’ activity and were further constrained within the boundaries of masks, from Stanford Willard Atlas(56) for MPFC and WFU PickAtlas(57) for all other regions. Pre-processed data underwent smoothing, and a generalized linear model was used to regress white matter and cerebrospinal fluid signals and 6 head motion parameters (3 translation and 3 rotational).

#### 3.3.4 Structural Connectivity Analyses

Group differences in structural connectivity were tested at two levels. First, edgewise connectivity was compared using two-sample t-tests across 28 unique undirected connections (forming the upper triangle of the symmetric structural connectivity matrix). Second, node strength (the sum of all connections for each region) was compared across eight network nodes. False discovery rate correction using the Benjamini– Hochberg method(58) was applied separately to edgewise and node-level tests. Effect sizes were reported as Cohen’s d.

#### 3.3.5 Hierarchical Empirical Bayes

Structurally informed effective connectivity analyses were conducted using spDCM(37,39) within a HEB framework(36), implemented in SPM12 using standard parametric empirical Bayes (PEB)(59) routines. Briefly, this approach models structurally informed resting-state effective connectivity at the group level, treating subject-level DCM parameters as data. The inferred structurally informed group-level effectivity connectivity and uncertainty is then propagated to the subject level by using the group level posterior as priors. At the subject level, spDCM was used to infer resting-state effective connectivity from fMRI data. This spDCM represents a spectral-domain transformation of a continuous-time state-space model, where neuronal states evolve according to effective connectivity encoded in a transition matrix, and are mapped to observed BOLD signals via a biophysical hemodynamic model(38). Next, we will describe this procedure in detail. For each subject, DCMs were inverted for a network of eight ROIs, yielding posterior estimates of effective connectivity. These subject-level posteriors were first entered into a HEB model with intercept-only design matrices and inverted under uninformed priors. At this stage, no covariates were included to establish a baseline model against which structurally informed priors could be evaluated.

##### 3.3.5.1 Establishing structure-based priors

After DCMs were inverted, subject-level posteriors were entered into HEB models—for all, no-depression and depression participants. Using intercept-only design matrix and uninformed priors, we then explored structure-based constraints *post hoc*. Specifically, we parameterised a linear mapping between structural connectivity and prior-variance (Fig. 1e), characterised by two hyperparameters: baseline variance (α) and scaling with structural connectivity (β). Varying these hyperparameters allowed us to evaluate a large model space of structure–function relationships. Bayesian model reduction was used to score models based on free energy, and Bayesian model averaging was applied to yield evidence-weighted transformations of prior variance (Fig. 2b–c). These transformations were compared between depression-status groups (Fig. 2c), and the associated gains in model evidence were quantified (Fig. 2d).

**Fig. 2.**
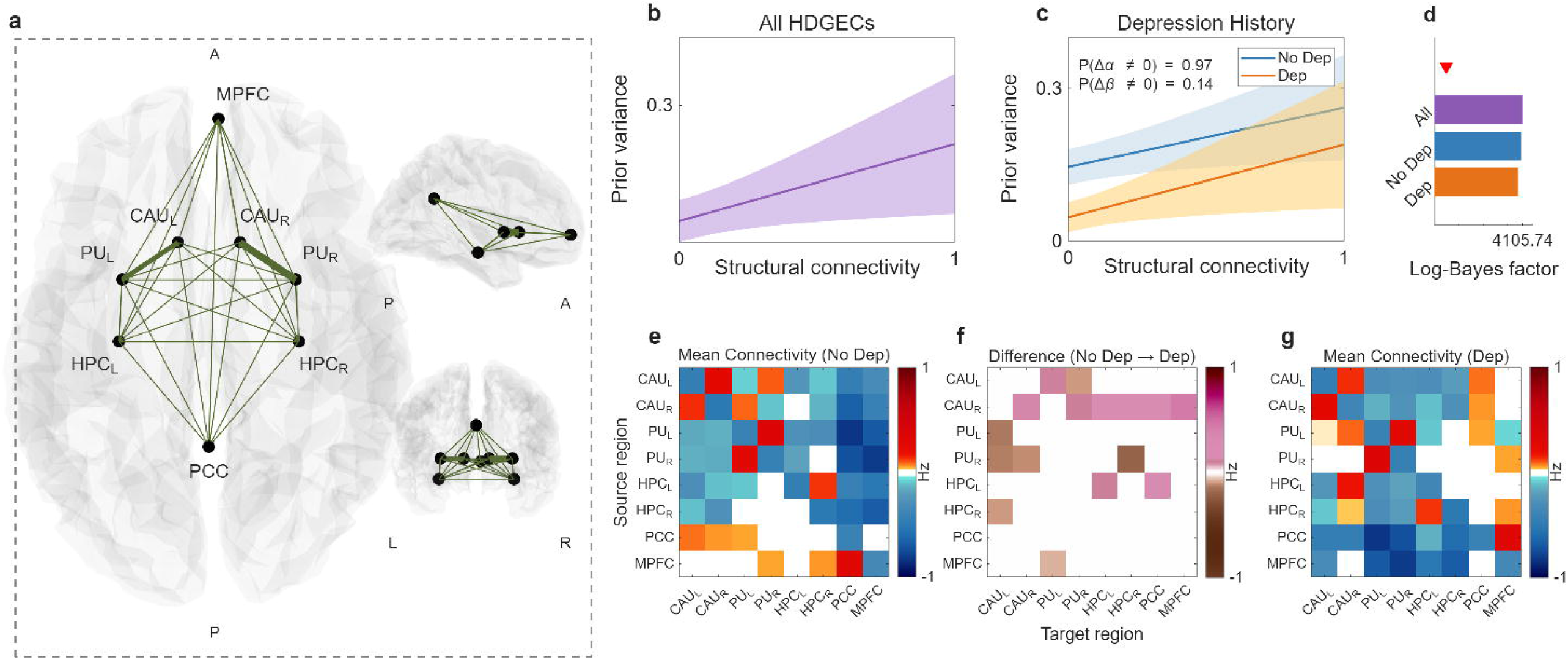
Structural connectivity constrains effective connectivity for HDGECs with depression history. **(a)** Schematic representation of default mode network (medial prefrontal cortex [MPFC], posterior cingulate [PCC], and bilateral hippocampi [HPC_L_, HPC_R_] and striatal (bilateral caudate [CAU_L_, CAU_R_] and putamen [PU_L_, PU_R_]) structural connectivity projected on axial, sagittal, and coronal planes. Line thickness indicates structural connection strength. **(b)** Model evidence-weighted prior-variance transformations show how structural connectivity scales *prior* variance of effective connectivity across all HDGECs. Shading denotes standard error envelope. **(c)** Depression history modulates the baseline variance (α) of effective connectivity, with HDGECs showing depression history (orange) versus those without (blue). P(Δα ≠ 0) and P(Δβ ≠ 0) indicate posterior probabilities (P) that group differences exist for α (variance) and β (slope) parameters, respectively. **(d)** Log-Bayes factors showing increased group-level model evidence. Red triangle indicates log-Bayes factor threshold of 3 (considered significant). Group-level mean effective connectivity patterns derived from hierarchical models incorporating depression status as a covariate: **(e)** HDGECs without depression history and **(g)**. HDGECs with depression history. Red indicates excitatory connections and blue indicates inhibitory connections. **(f)** Between-group differences in effective connectivity comparing HDGECs with depression history versus those without, derived from covariate models. Purple indicates increasing connectivity while brown indicates decreasing connectivity. Off-diagonal entries represent directed inter-regional influences in (Hz). Positive evidence set at 75% posterior probability throughout.

The resulting posterior estimates were integrated within a three-level hierarchical Bayesian framework. At the first level, fMRI time series were generated from subject-specific effective connectivity. At the second level, subject-level connectivity parameters were modelled as functions of group effects (e.g., depression status), with random effects capturing inter-individual variability. At the third level, group-level connectivity parameters were modelled as deviations from hierarchical prior expectations, enabling inference about population-level network organizationunder either uninformed or structurally informed priors. Mathematical details are provided in supplementary methods.

##### 3.3.5.2 Group-differences in structurally informed effective connectivity

Finally, to examine depression-related effects, we specified design matrices encoding history of depression, and covariates (sex, mood-related medication use, and scanner effects [SI.Fig, 1]). Group-level analyses examined effective connectivity across the eight DMN and striatal ROIs, testing for group-level differences in effective connectivity under structural constraints. Posterior probabilities were thresholded at Pp ≥ 0.75, consistent with positive evidence. To assess associations between differences in effective connectivity and clinically elevated HADS-D or BDI-II scores versus non-elevated, analyses were constrained to only significant connections and the structural connectivity priors from the difference analysis were used, alongside the same covariates.

To test predictive validity, we conducted leave-one-out cross-validation (56). For each held-out participant,(60) connectivity estimates were predicted from models fit to the remaining sample. Analyses focused on connections that showed significant group differences, and predictions were compared to depression severity cur-offs (HADS-D and BDI-II). Prediction accuracy was assessed using Pearson’s correlations.

### 3.4 Data availability

No new data were collected. The data that support the findings of this study are available on the Enroll-HD platform via request at https://www.enroll-hd.org/for-researchers/access-data-biosamples/.

## 4 Results

Analyses included 98 premanifest HDGECs (history of depression: *n* = 30, *M*_age_ = 43.13 years, *SD* = 6.68, 63.33% female, *M*_CAG_ = 43.10, *SD* = 1.90; no history of depression: *n* = 68, *M*_age_ = 42.68 years, *SD* = 10.32, 42.65% female, *M*_CAG_ = 43.43, *SD* = 2.43) from TrackOn-HD (12,22) as previously reported(18).

### 4.1 Structural Connectivity

We found no differences in edge or node strength in any ROIs between HDGECs with a depression history compared to those without (S.Table 4–5). Therefore, a group-level structural connectome across both groups was used for HEB analyses.

### 4.2 Structure–function dynamics in depression for HDGECs

Fig. 2a shows group average structural connectivity for the DMN and striatum (line weight is proportional to connection strength after correcting for spatial and geometric confounds).

The evidence-weighted prior-variance transformation analysis showed that structural connectivity influenced effective connectivity patterns across all HDGECs, with stronger structural connections allowing effective connections to be more variable (i.e. stronger) as well (Fig. 2b-c). Between-group comparison of model evidence-weighted hyperparameters revealed an intercept shift (α) in HDGECs with depression history, where the depression group showed lower baseline variability in effective connectivity, such that near-zero-valued structural connections—the MPFC–left-hippocampus connection being the nearest zero—resulted in effective connections being more tightly regularized toward zero (Fig. 2c). Group-level model evidence substantially exceeded the conventional log-Bayes factor threshold of 3(61)(Fig. 2d), providing strong statistical support for incorporating structural connectivity priors in effective connectivity estimation.

HEB models incorporating structural connectivity-informed priors were subsequently inverted to characterize group-specific effective connectivity architectures. Maximum *a posteriori* estimates showed distinct connectivity profiles for HDGECs without depression history (Fig. 2e) and those with depression history (Fig. 2g), alongside depression-related connectivity alterations (Fig. 2f). Both groups exhibited predominantly inhibitory default mode network connectivity patterns, coupled with mixed excitatory and inhibitory striatal connections. Connections from DMN to striatum were excitatory for HDGECs without depression and inhibitory for those with a history of depression, and this was reversed for connections from striatum to DMN. When looking at differences in the strength of connectivity, HDGECs with depression history demonstrated reduced inhibitory self-connectivity within the right caudate nucleus, increasing right caudate efferents, and increasing inhibitory control of DMN on striatum.

### 4.3 Depression symptom severity-associated effective connectivity

Depression severity measures revealed distinct connectivity association patterns between groups (Fig. 3a-l).

**Fig. 3.**
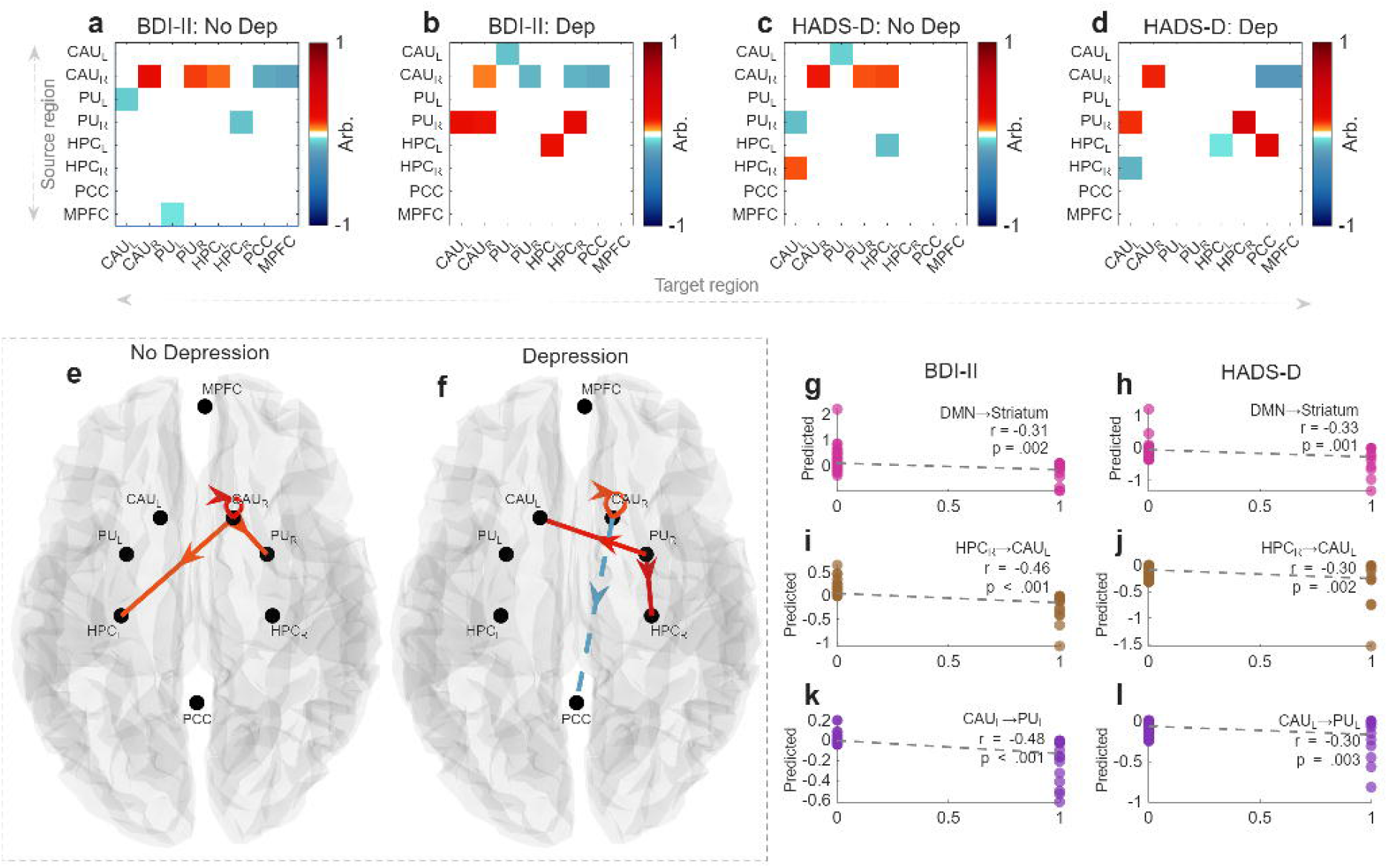
Depression severity associations and predictive connectivity patterns by depression history. Effective connectivity associations with depression severity (meeting clinical cut-offs for elevated symptoms) by depression history group. **(a-d)** Connectivity matrices showing associations between effective connectivity and depression severity for Beck depression inventory, 2nd edition (BDI-II) in participants **(a)** without depression history and **(b)** with depression history, and hospital anxiety and depression scale, depression subscale (HADS-D) in participants **(c)** without depression history and **(d)** with depression history. Red indicates positive associations, and blue indicates negative associations with depression severity (posterior probability ≥ 75%). **(e-f)** Brain visualizations of significant connectivity patterns across symptom measures for participants **(e)** without depression history and **(f)** with depression history. **(g-l)** Leave-one-out cross-validation correlations between predicted and actual depression scores for specific connections: **(g-h)** DMN to striatum connections (right hippocampus to left caudate and medial prefrontal cortex to left putamen) for BDI-II and HADS-D **(i-j)** right hippocampus to left caudate for BDI-II and HADS-D, and **(k-l)** left caudate to left putamen connections for BDI-II and HADS-D. Correlation coefficients (*r*) and p-values are displayed. CAU = caudate; PU = putamen; HPC = hippocampus; PCC = posterior cingulate cortex; MPFC = medial prefrontal cortex; DMN = default mode network.

In HDGECs without depression history, both BDI-II (Fig. 3a) and HADS-D (Fig. 3c) scores demonstrated associations with effective connectivity across default mode and striatal regions. The no depression history group showed positive right caudate efferents, including self-connectivity, and to both right putamen and left hippocampus (Fig. 3e). HDGECs with depression history exhibited different association patterns for both BDI-II (Fig. 3b) and HADS-D (Fig. 3d). Common connectivity associations also included positive associations with right caudate self-connectivity but differed with negative association in efferents to PCC. Right putamen efferents to left caudate and right hippocampus were also positively associated with depression severity. (Fig. 3f).

Leave-one-out cross-validation assessed the predictive utility of specific connections that showed significant associations with depression severity. DMN to striatum connections (i.e., right hippocampus to left caudate and MPFC to left putamen) demonstrated significant predictive capacity for both clinically significant BDI-II and HADS-D scores (Fig. 3g-h). Right hippocampus to left caudate (Fig. 3i-j) and left caudate to left putamen connections (Fig. 3k-l) predicted both BDI-II and HADS-D clinically elevated symptoms. Additional individual connections that met significance for only one depression symptom measure are shown in SI.Fig 2.

## 5 Discussion

We applied a novel HEB framework with structurally informed priors to assess two competing theoretical frameworks for depression-related network reorganization: functional constraint where neural communication becomes increasingly determined by underlying anatomical architecture, versus functional decoupling, where brain networks maintain independence from structural limitations through pathological mechanisms. Our findings provide evidence for reduced network flexibility, particularly for weakly connected pathways, in depression: HDGECs with depression history showed lower baseline variability in effective connectivity (reduced α parameter), such that near-zero-valued structural connections became more tightly regularized toward zero. The potentially reflects an early marker of network dysfunction in depression for premanifest HDGECs.

First, we demonstrate that incorporating structural information into spDCM via HEB procedures resulted in increased model evidence for all HDGECs, showing that anatomical architecture provides informative constraints that improve the plausibility of inferred effective connectivity. Second, our findings extend prior work in Alzheimer’s disease, where correlation-based approaches have reported both structure–function decoupling(28) and increased coupling within DMN and rich-club networks(30), but without mechanistic resolution. We move beyond correlations to show how anatomical priors constrain functional dynamics in HDGECs with depression history. Increased structure–function coupling within the DMN has also been reported in major depression(32), consistent with our observation that depression history in HDGECs is associated with an intercept shift reflecting a global reduction in variance across connections. This pattern indicates that effective connectivity becomes more rigidly governed by structural architecture, with reduced flexibility for functional interactions, particularly for connections with minimal structural connectivity. The intercept shift specifically increased regularization of near-zero-valued structural connections (such as MPFC-hippocampus) also toward zero-valued effective connectivity, while leaving strongly connected pathways relatively unaffected. Notably, these findings were obtained in the same HDGEC cohort previously analyzed with predictive modelling, which revealed compensatory functional connectivity in regions with weaker structural connectivity(27). By contrast, our mechanistic framework shows that in the presence of depression, functional interactions are no longer flexibly upregulated but instead compressed into a narrower variance regime, becoming more deterministically governed by anatomy. While correlation-based analyses capture adaptive functional adjustments to structural deficits, HEB modelling reveals an overarching constraint: in HDGECs with depression, structural architecture increasingly dominates functional dynamics, limiting the scope for compensation.

Extending prior work linking depression in HDGECs to increased anterior–posterior DMN functional connectivity and reduced DMN–striatal structural connectivity(17), our structurally-informed results isolate directional pathways through which depression reshapes networks. We found the valence of effective connectivity largely agreed with previous structurally uninformed effective connectivity models(18) revealing consistent within striatal connectivity patterns (e.g., right to left caudate excitation, left to left putamen inhibition, and left caudate self-inhibition), and recapitulating inhibitory hippocampal afferents to striatal regions—patterns that align with hippocampal dysfunction in non-neurological major depression (62,63). In contrast, structurally uninformed low-magnitude excitatory efferent PCC and MPFC connections in uninformed models(18) manifested as inhibitory in the present anatomically constrained analyses for HDGECs with depression history. Despite being a neural marker of major depression, we observed no influence of MPFC on PCC(64), consistent with previous structurally uninformed effective connectivity models(18).

Further, depression-related alterations demonstrate convergence with spDCM(18) through increased connectivity from efferent right caudate, while anatomically constrained modeling revealed minimal DMN influence outside of left hippocampal efferents. Lateralized connectivity alterations of striatum complement broader evidence for right hemisphere compensation in HDGECs(21), where increased functional connectivity between preserved regions represents adaptive network reorganization(22). Effective connectivity associations with clinically significant depression symptoms revealed that both structurally informed and uninformed approaches(18) identified right caudate self-connectivity as a marker of depression symptom severity, exhibiting positive correlations with meeting clinical cut-offs in HDGECs with and without depression history in the present study. Findings posit striatal-hippocampal circuitry as potential hubs for depression severity, suggesting effective connectivity alterations reflect functional reorganization mechanisms responding to underlying structural vulnerabilities in the striatum.

The present cross-sectional analysis provides foundation for several future directions. Firstly, longitudinal HEB implementations could establish clinical trajectories by tracking the evolution of structure–function mapping parameters. While intercept shifts capture baseline variance dampening magnitude, slope changes would reflect temporal variations in the rate of constraint between functional and structural architectures. Should steepening slopes be observed longitudinally, this could indicate progressive functional network collapse where effective connectivity becomes increasingly constrained toward structural architecture, potentially reflecting advancing neurodegeneration that reduces capacity for functional reorganization independent of anatomical constraints. The direction and magnitude of such longitudinal changes in HDGECs remains an empirical question requiring future investigation. Following this, whole-brain spectral DCM which is scaled up by using faster implementations(65), would allow for detection of additional depression-relevant circuits. For example, inclusion of the cognitive control network (prefrontal cortex, ventral striatum, dorsal striatum) linked to anhedonia and the affective network (orbitofrontal cortex, anterior cingulate cortex, limbic regions) linked to dysphoria(66). Aligning circuitry with unique aspects of HD depression in which canonical depressive symptoms (low mood, emotional lability, anhedonia, hopelessness, guilt) co-occur with distinctive features such as anger and irritability(67), would further our understanding of the pathophysiology of depression in HDGECs. As HD therapeutic trials develop, structure–function dynamics warrants investigation as a potential neuroimaging biomarker(68), with future longitudinal studies needed to establish whether structure–function dynamics could stratify participants by degree of functional network constraint or if longitudinal slope tracking could provide indices of therapeutic effects on network flexibility.

In conclusion, depression systematically reduced network flexibility in premanifest HDGECs through variance dampening of effective connectivity, constraining functional networks toward their structural architecture. Convergence between structurally informed and uninformed approaches revealed consistent depression signatures, with anatomical constraints unmasking inhibitory projections of DMN hub regions that appeared excitatory in unconstrained analyses. Effects centered on striatal and hippocampal efferents, with right caudate self-connectivity and altered hippocampal-putamen interactions that associated with depression severity. Structurally informed effective connectivity provides a mechanistic framework for understanding how depression emerges for premanifest HDGECs.

## 6 Supplementary material

Supplementary material is available online.

## 7 Author contributions

TB, MDG and AR conceived the study. TB and MDG developed the methodology, implemented the software, performed formal analyses and investigations, curated data, and contributed to visualization and project administration. TB additionally drafted the original manuscript. LN contributed to methodology development. AR provided supervision and funding acquisition. All authors (TB, MDG, LN, YGJ, AR) contributed to the review and editing of the manuscript.

