## Supplementary Materials for "Depression reduces structurally informed network flexibility in premanifest Huntington’s disease"

3 Wellcome Centre for Human Neuroimaging, University College London, London, United Kingdom

### Methods

#### Clinical measures

Four HD-related measures were included. The Unified Huntington’s Disease Rating Scale (UHDRS) Total Motor Score (TMS) is a 15-item clinician-rated severity of motor signs with a maximum score of 124(1). The CAG-Age Product (CAP) score models the interaction of age and CAG repeat length on disease progression where the score is 100 at expected age of diagnosis(2–5). The Disease Burden Score (DBS) indexes exposure to huntingtin by modelling age and CAG repeat, and includes a constant of 35.5(4). The HD Integrated Staging System (HD-ISS) incorporates pathophysiology alongside clinical and functional changes, and includes stages 0–3 based on genetic confirmation, striatal volume changes, cognitive impairment, and functional decline(6).

The Beck Depression Inventory, 2nd Edition (BDI-II; 7) has 21 items on a 4-point Likert scale (maximum score of 63) assessing depressive symptoms over a two-week period(7). The BDI-II has been recommended in HD(8) and a cut-off score of 10/11 discriminates clinically elevated depression in HD, with sensitivity of 1.00 and specificity of 0.66(9). The Hospital Anxiety and Depression Scale, Depression Subscale (HADS-D; 10) has seven items on a 4-point Likert scale, with a maximum score of 21(10). Research in HD suggests a cut-off of 6/7 discriminates clinically elevated depression, with sensitivity of 1.00, and specificity of 0.82(9).

#### MRI acquisition

3T MRI data was acquired on two different scanner systems: Philips Achieva at Leiden and Vancouver and Siemens TIM Trio at London and Paris, described in detail elsewhere(11). For T1-weighted image acquisition, a 3D MPRAGE sequence was used with the following parameters: TR 2200ms (Siemens)/7.7ms (Philips), TE 2.2ms (S)/3.5ms (P), FOV 280mm (S)/240mm (P), flip angle 10°(S)/8°(P), 208(S)/164(P) sagittal slices (slice thickness: 1.1mm, gap: no gap, matrix size 256×256 (S)/224×224 (P)) and bandwidth of 240Hz (S)/241Hz (P) per participant(12). For resting-state fMRI, whole-brain volumes were acquired using a T2*-weighted echo planar imaging (EPI) sequence with the following parameters: TR 3000ms, TE 30ms, FOV 212mm, flip angle 80°, 48 slices in ascending order (slice thickness: 2.8mm, gap: 1.5mm, in-plane resolution 3×3×3mm) and bandwidth of 1906Hz per pixel. Rs-fMRI data were collected first, then both sets of task fMRI data, with 165 volumes acquired per participant (Klöppel et al., 2015). For diffusion-weighted imaging, whole-brain volumes were acquired using a single-shot echo-planar imaging (EPI) sequence with the following parameters: TR 13,100ms (Siemens)/11,193ms (Philips), TE 88ms (S)/56ms (P), FOV 256mm, flip angle 90°, 72 axial slices in interleaved order (slice thickness: 2.0mm, no gap, in-plane resolution 2.0×2.0mm) with phase encoding in opposite directions between scanner types (posterior-anterior (S)/anterior-posterior (P)). Diffusion-weighted images were acquired with 41 unique gradient directions (b = 1000 s/mm²) and seven images with no diffusion weighting (b = 0 s/mm²). The acquisition utilized parallel imaging acceleration with GRAPPA (S)/SENSE (P) and partial Fourier factors of 0.75 (S)/0.54 (P). Total scanning time was approximately 12 minutes for T1-weighted, 10 minutes for diffusion-weighted imaging, and 15 minutes for resting-state fMRI(11).

#### Preprocessing

Firstly data was reorganised to BIDS specification to standardise the data structure, ensure dataset integrity, and define the readable metadata(13). Preprocessing was performed for all participants using *fMRIPrep* 21.0.2(14,15; RRID:SCR_016216)—described in detail below—and MRIQC v22.0.6(16), which both used FreeSurfer v-6.0.1(17). Spatial smoothing by a 6 mm full-width half-maximum Gaussian kernel was subsequently undertaken using SPM12 (7771). MRIQC is a quality control tool for anatomical and fMRI data that extracts no-reference image quality metrics for scan quality assessments(16). It complimented fMRIPrep and provided group level quality control assessments. Preprocessed derivatives from fMRIPrep provided the input for subsequent DCM analyses.

##### fMRIPrep

Results included in this manuscript come from preprocessing performed using *fMRIPrep* 21.0.2(18,19; RRID:SCR_016216), which is based on *Nipype* 1.6.1(20,21; RRID:SCR_002502).

###### Anatomical data preprocessing

A total of 3 T1-weighted (T1w) images were found within the input BIDS dataset. All of them were corrected for intensity non-uniformity (INU) with N4BiasFieldCorrection(22), distributed with ANTs 2.3.3(23; RRID:SCR_004757). The T1w-reference was then skull-stripped with a *Nipype* implementation of the antsBrainExtraction.sh workflow (from ANTs), using OASIS30ANTs as target template. Brain tissue segmentation of cerebrospinal fluid (CSF), white-matter (WM) and gray-matter (GM) was performed on the brain-extracted T1w using fast (FSL 6.0.5.1:57b01774, RRID:SCR_002823; 24). A T1w-reference map was computed after registration of 3 T1w images (after INU-correction) using mri_R_obust_template (FreeSurfer 6.0.1; 25). Brain surfaces were reconstructed using recon-all (FreeSurfer 6.0.1, RRID:SCR_001847; 26), and the brain mask estimated previously was refined with a custom variation of the method to reconcile ANTs-derived and FreeSurfer-derived segmentations of the cortical gray-matter of Mindboggle (RRID:SCR_002438; 27). Volume-based spatial normalization to one standard space (MNI152NLin2009cAsym) was performed through nonlinear registration with antsRegistration (ANTs 2.3.3), using brain-extracted versions of both T1w reference and the T1w template. The following template was selected for spatial normalization: *ICBM 152 Nonlinear Asymmetrical template version 2009c* [RRID:SCR_008796; TemplateFlow ID: MNI152NLin2009cAsym](28).

###### Functional data preprocessing

For each of the 3 BOLD runs found per subject (across all tasks and sessions), the following preprocessing was performed. First, a reference volume and its skull-stripped version were generated using a custom methodology of *fMRIPrep*. Head-motion parameters with respect to the BOLD reference (transformation matrices, and six corresponding rotation and translation parameters) are estimated before any spatiotemporal filtering using mcflirt (FSL 6.0.5.1:57b01774; 29). BOLD runs were slice-time corrected to 1.48s (0.5 of slice acquisition range 0s-2.95s) using 3dTshift from AFNI (30; RRID:SCR_005927). The BOLD time-series (including slice-timing correction when applied) were resampled onto their original, native space by applying the transforms to correct for head-motion. These resampled BOLD time-series will be referred to as *preprocessed BOLD in original space*, or just *preprocessed BOLD*. The BOLD reference was then co-registered to the T1w reference using bbregister (FreeSurfer) which implements boundary-based registration (31). Co-registration was configured with six degrees of freedom. Several confounding time-series were calculated based on the *preprocessed BOLD*: framewise displacement (FD), DVARS and three region-wise global signals. FD was computed using two formulations following Power (absolute sum of relative motions (32), and Jenkinson (relative root mean square displacement between affines (29)). FD and DVARS are calculated for each functional run, both using their implementations in *Nipype* (following the definitions by Power et al. 2014). The three global signals are extracted within the CSF, the WM, and the whole-brain masks. Additionally, a set of physiological regressors were extracted to allow for component-based noise correction (*CompCor*; 33)). Principal components are estimated after high-pass filtering the *preprocessed BOLD* time-series (using a discrete cosine filter with 128s cut-off) for the two *CompCor* variants: temporal (tCompCor) and anatomical (aCompCor). tCompCor components are then calculated from the top 2% variable voxels within the brain mask. For aCompCor, three probabilistic masks (CSF, WM and combined CSF+WM) are generated in anatomical space. The implementation differs from that of Behzadi et al. in that instead of eroding the masks by 2 pixels on BOLD space, the aCompCor masks are subtracted a mask of pixels that likely contain a volume fraction of GM. This mask is obtained by dilating a GM mask extracted from the FreeSurfer’s *aseg* segmentation, and it ensures components are not extracted from voxels containing a minimal fraction of GM. Finally, these masks are resampled into BOLD space and binarized by thresholding at 0.99 (as in the original implementation). Components are also calculated separately within the WM and CSF masks. For each CompCor decomposition, the *k* components with the largest singular values are retained, such that the retained components’ time series are sufficient to explain 50 percent of variance across the nuisance mask (CSF, WM, combined, or temporal). The remaining components are dropped from consideration. The head-motion estimates calculated in the correction step were also placed within the corresponding confounds file. The confound time series derived from head motion estimates and global signals were expanded with the inclusion of temporal derivatives and quadratic terms for each(34). Frames that exceeded a threshold of 0.5 mm FD or 1.5 standardised DVARS were annotated as motion outliers. The BOLD time-series were resampled into standard space, generating a *preprocessed BOLD run in MNI152NLin2009cAsym space*. First, a reference volume and its skull-stripped version were generated using a custom methodology of *fMRIPrep*. All resamplings can be performed with *a single interpolation step* by composing all the pertinent transformations (i.e. head-motion transform matrices, susceptibility distortion correction when available, and co-registrations to anatomical and output spaces). Gridded (volumetric) resamplings were performed using antsApplyTransforms (ANTs), configured with Lanczos interpolation to minimize the smoothing effects of other kernels(Lanczos 1964). Non-gridded (surface) resamplings were performed using mri_vol2surf (FreeSurfer).

Many internal operations of *fMRIPrep* use *Nilearn* 0.8.1(35; RRID:SCR_001362), mostly within the functional processing workflow. For more details of the pipeline, see [the section corresponding to workflows in *fMRIPrep*’s documentation](https://fmriprep.readthedocs.io/en/latest/workflows.html).

###### Copyright waiver

The above boilerplate text was automatically generated by fMRIPrep with the express intention that users should copy and paste this text into their manuscripts *unchanged*. It is released under the [CC0](https://creativecommons.org/publicdomain/zero/1.0/) license.

##### Diffusion MRI preprocessing

Raw data were converted to MRtrix3 format and T1 images underwent reorientation, field-of-view cropping, and brain extraction (fractional intensity threshold = 0.3). Preprocessing comprised sequential denoising using random matrix theory(36), Gibbs ringing artifact removal(37) and motion and distortion correction. Given the absence of reverse phase-encoding acquisitions, susceptibility distortions were corrected using Synb0-DisCo (v3.1; 38). Zero-padding was applied, then motion, eddy current, and susceptibility distortion correction was performed with slice-to-volume correction and outlier replacement(39–41). Brain masks were generated (fractional intensity threshold = 0.2; 42) followed by bias field correction using the N4 algorithm(22). Following preprocessing, systematic quality assurance was performed by two independent researchers. Quality assurance included assessment of eddy current outlier detection and visual inspection of distortion correction and brain mask accuracy. Brain extraction parameters were iteratively adjusted until inter-rater concordance was achieved.

White matter fiber orientation distributions were estimated using multi-shell multi-tissue constrained spherical deconvolution with the dhollander algorithm for response function estimation(43,44). Multi-tissue intensity normalization was applied across white matter, gray matter, and cerebrospinal fluid following three-tissue CSD modelling(43,45). Anatomically-constrained tractography was performed using the iFOD2 algorithm, generating 4 million streamlines from gray-white matter interface seeds(46,47). Five-tissue-type segmentation masks were created from T1 images using automated segmentation(24,48). Streamline weights were optimized using SIFT2 to ensure biologically plausible tract densities(49).

Whole-brain structural connectivity matrices were constructed using a custom parcellation. Atlas registration employed a two-stage approach: brain-extracted T1 images were registered to MNI152 space using ANTs symmetric normalization(23), followed by transformation to native DWI space via linear registration(29,50). Connectivity matrices were generated using streamline counts with inverse node volume scaling and symmetric averaging, followed by label conversion to standardized region-of-interest definitions using MRtrix3.

#### Statistical analyses

##### Hierarchical empirical bayes

###### First level

Effective connectivity was inferred using a hierarchical empirical Bayes framework built on the well-validated spectral DCM for resting-state fMRI(51–53), and standard routines implemented in the SPM toolbox(54). Full details of the generative model, prior specification, and inversion procedure are reported in previous work(55). Here, we provide a high-level summary.

Starting at the first (subject) level, for $s=1,\ldots,S$ subjects, we consider a DCM that describes how $i=1,\ldots,n$ neuronal populations influence one another and give rise to observed BOLD responses. This DCM is inverted—using variational Bayes under the Laplace approximation(56)—in the spectral domain(57), yet assumes—in the time domain—the following state-space representation:

|  | $\dot{\boldsymbol{x}}(t)=\boldsymbol{Ax}(t)+\boldsymbol{v}(t)$ (*state equation*)  $\hat{\boldsymbol{y}}\left( t \right)=h\left( \boldsymbol{x}\left( t \right),\boldsymbol{\theta}_{h} \right)+\boldsymbol{e}(t)$ (*observation equation*). | (1) |
| --- | --- | --- |

Here, $\boldsymbol{x}\left( t \right)\boldsymbol{=}\left[ x_{1}(t), \ldots,x_{n}(t) \right]^{T}$ denotes the neuronal state vector, where each scalar function $x_{i}(t\mathbb{)\in R}$ describes the ensemble activity of the $i$-th population at time $t$. The transition matrix $\boldsymbol{A}\in\mathbb{R}^{n\times n}$ encodes the intra- and inter-regional modulation of the rates of change in this ensemble activity (the effective connectivity), in its diagonal and off-diagonal elements, respectively. In the observation equation, $h$ is the hemodynamic response function (HRF) with free parameters $\boldsymbol{\theta}_{h}$, which maps ensemble neuronal activity to expected BOLD responses $\hat{\boldsymbol{y}}\left( t \right)$, which are compared to empirical data with parameterized residual precision. Both endogenous fluctuations $\boldsymbol{v}(t)$, and observation error $\boldsymbol{e}(t)$, are parameterized as power-law noise.

###### Second level

With DCMs inverted for a specified network of regions, effective connectivity posterior means are moved into the hierarchical empirical Bayes model. Let $\boldsymbol{\theta}^{\left( 1 \right)}$ be a vector of stacked posterior means for $S$ subjects, such that $\boldsymbol{\theta}^{\left( 1 \right)}=\left[ \mathrm{vec}\left( \boldsymbol{\theta}_{1}^{\left( 1 \right)} \right);\ldots;\mathrm{vec}\left( \boldsymbol{\theta}_{S}^{\left( 1 \right)} \right) \right]\in\mathbb{R}^{Sp}$. Here, $\boldsymbol{\theta}_{s}^{\left( 1 \right)}=\boldsymbol{A}_{s}$ are the effective connectivity parameters for the $s$-th subject, and $p=n^{2}$ is the number of the $s$-th subject’s parameters considered. We can now consider a hierarchical regression formulation

|  | $\boldsymbol{\theta}^{\left( 2 \right)}\mathcal{=N}\left( \boldsymbol{\mu}^{\left( 3 \right)},\boldsymbol{\Sigma}^{(3)} \right)$ (*third-level model*)  $\boldsymbol{\theta}^{\left( 1 \right)}=\left( \boldsymbol{X\bigotimes I} \right)\boldsymbol{\theta}^{\left( 2 \right)}+\mathcal{N}\left( \mathbf{0}\boldsymbol{,}\boldsymbol{\Sigma}^{(2)} \right)$ (*second-level model*) | (2) |
| --- | --- | --- |

Here, $\boldsymbol{\theta}^{\left( 2 \right)}\in\mathbb{R}^{pq}$ is the group-level effective connectivity, $q$ is the number of covariates (including the intercept) encoded in a second-level design matrix $\boldsymbol{X\in}\mathbb{R}^{S\times q}$, and the Kronecker product $\boldsymbol{X\bigotimes I\in}\mathbb{R}^{Sp\times pq}$—where $\boldsymbol{I}\in\mathbb{R}^{p\times p}$ is the identity matrix—ensures that both $\boldsymbol{\theta}^{\left( 2 \right)}$ and the subject-specific random effects $\mathcal{N}\left( \mathbf{0}\boldsymbol{,}\boldsymbol{\Sigma}^{(2)} \right)$, are appropriately tiled across parameters $\boldsymbol{\theta}^{\left( 1 \right)}$. The second-level covariance $\boldsymbol{\Sigma}^{(2)}$ is parametrized in terms of scaled precision components(58), whereas the third-level covariance $\boldsymbol{\Sigma}^{(3)}$ incorporates structure-based constraints.

In the single-intercept case—$q=1$, $\boldsymbol{X=}\boldsymbol{1}_{S\times1}$—the third-level covariance $\boldsymbol{\Sigma}^{(3)}$, is specified such that variance for between-region connections scales with normalized structural connectivity $\tilde{\boldsymbol{C}}\in\left[ 0,1 \right]^{n\times n}$, according to two hyperparameters: $\alpha$ sets a baseline variance, and $\beta$ controls how strongly variance increases with structural connectivity strength. Self-connections are assigned a fixed small variance $\delta=1/64$. Formally:

$$\boldsymbol{\Sigma}^{(3)}\boldsymbol{=}\mathrm{diag}\left( \mathrm{vec}\left( \delta\boldsymbol{I+}\left( \alpha\boldsymbol{11}^{\boldsymbol{T}}\boldsymbol{+}\beta\tilde{\boldsymbol{C}} \right)\circ\left( 1-\boldsymbol{I} \right) \right) \right)$$

(3)

Here, $\boldsymbol{11}^{\boldsymbol{T}}$ is a matrix of ones, $\boldsymbol{I}$ is the $n\times n$ identity matrix, and $\circ$ is the elementwise product. In all cases where $q>1$, $\boldsymbol{\Sigma}^{(3)}$ was extended per the method described by Zeidman and colleagues(58).

###### Procedures

To obtain evidence-weighted hyperparameters, we first inverted uniformed models—the case where $q=1,$ $\alpha=1/2$ and $\beta=0$—and then explored (reduced) structurally informed models nested within (full) uninformed models by performing a grid search over values of $\alpha$ and $\beta$, and using Bayesian model reduction to score structurally informed models in terms of their free energy (a lower-bound of the model evidence; 54). Evidence-weighted hyperparameters were then obtained by taking the mean across the hyperparameter grid, weighted by a softmax transformation of the model free energies(59). This is how we obtained the evidence-weighted prior-variance transformations for all subjects, and depression history and no-depression history subgroups (Fig. 1). In all subsequent analyses, we used the evidence-weighted prior-variance transformation obtained for all subjects (Fig. 1b), to obtain structure-based priors (Fig. 2e–g and Fig. 3).

Once evidence-weighted hyperparameters were established, hierarchical empirical Bayes models were used for two purposes. First, to assess whether structurally informed priors improved model evidence, we specified second-level design matrices ($\boldsymbol{X}$) with an intercept (column 1) and covariates for sex, scanner site, and medication use (columns 2–4), but no depression or clinical symptom severity regressor (Fig. 1b-c).

To test for differences in effective connectivity for HDGEC with and without a depression history (Fig. 2e–g) the design matrix was extended to include the group/clinical regressor (column 2) in addition to the and same covariates (which were z-scored). Posterior parameter estimates were thresholded at posterior probability ≥ 0.75 (for positive evidence) and thus provided effective connectivity estimation (valence and strength) and regression coefficient associations with clinical measures.

To test effective connectivity associations with clinically elevated depression symptoms (BDI-II, HADS-D; Fig. 3a-f), we applied the evidence-weighted structural priors derived from the group difference analysis (depression history) to constrain the prior covariance matrix, informing prior variances for significant effective connections from the difference analysis (depression history) while setting uninformative priors for connections below the significance threshold. The design matrix used the same covariate setup as the group difference analysis, with behavioral regressors for depression symptom severity replacing the group regressor, and all covariates were z-scored. Analyses were run separately for each depression history group and both symptom severity cut-offs, resulting in four analyses. Posterior parameter estimates were thresholded at posterior probability ≥ 0.75.

For all HEB models, the third-level prior covariance was derived from group-specific covariance matrices: variance terms from the no-depression group (either no depression history or no clinically elevated symptoms) informed the intercept, while the group regressor (when present) combined variance terms from both groups. Covariates were given uninformative priors to allow adjustment without constraining their effects, and before inversion the covariance matrix was rescaled to ensure design-scale invariance.

###### Self-connectivity in dynamic causal modelling

Within the DCM framework, self-connections are modelled as inhibitory to prevent potential runaway excitation; however, these self-connection parameters are logarithmically scaled (using the transformation log(−2*a*) in SPM). This logarithmic scaling is employed to enhance the numerical stability of the model fitting procedures and is technically motivated by using log-normal priors to enforce recurrent self-inhibition. Consequently, these self-connections can take both positive and negative values, with specific interpretations.

In SPM's reporting convention, a zero value (arbitrarily set) for a self-connection corresponds to −0.5 Hz, representing the default prior self-connectivity value. A positive self-connection signifies a relative increase in inhibition (faster decay rates, below 0.5 Hz), while a negative self-connection indicates a relative decrease in inhibition (slower decay rates in the −0.5 to 0 Hz range). These inhibitory self-connections regulate the gain or sensitivity to inputs from other regions. Decreased self-inhibition implies increased synaptic gain or sensitivity to inputs, whereas increased self-inhibition suggests a reduction in synaptic gain or sensitivity to inputs. Importantly, only self-connections are subjected to this logarithmic scaling transformation in DCM.

### Results

#### Participant characteristics

HD gene expansion carriers (HDGECs) did not differ in sex, age, ethnicity, handedness, or HD-related variables. HDGECs with a history of depression had significantly higher history of suicidal ideation, use of mood medication and HADS-D scores. Demographic variables were compared using Pearson’s chi-squared test or linear model ANOVA. Assumptions were met. Some variables demonstrated non-normal distributions: this was considered acceptable as it did not impact HEB analyses, and have been previously reported in full(60). Participant characteristics are summarized in SI.Table 1.

**SI.Table 1 Demographic, clinical, and psychiatric characteristics**

|  | **History of depression (N=30)** | **No history of depression (N=68)** | **p value** |
| --- | --- | --- | --- |
| Sex |  |  | 0.059^a^ |
| Female | 19 (63.33%) | 29 (42.65%) |  |
| Male | 11 (36.67%) | 39 (57.35%) |  |
| Age |  |  | 0.824^b^ |
| Mean (SD) | 43.13 (6.68) | 42.68 (10.32) |  |
| Range | 28.00 - 60.00 | 23.00 - 68.00 |  |
| Ethnicity |  |  | 0.639^a^ |
| American - Latin | 1 (3.33%) | 1 (1.47%) |  |
| Asian - east | 0 (0.00%) | 2 (2.94%) |  |
| Mixed | 0 (0.00%) | 1 (1.47%) |  |
| White | 29 (96.67%) | 64 (94.12%) |  |
| Handedness |  |  | 0.089^a^ |
| Ambidextrous | 2 (6.67%) | 2 (2.94%) |  |
| Left-handed | 4 (13.33%) | 2 (2.94%) |  |
| Right-handed | 24 (80.00%) | 64 (94.12%) |  |
| **HD-related Variables** | | | |
| CAG repeat length |  |  | 0.515^b^ |
| Mean (SD) | 43.10 (1.90) | 43.43 (2.43) |  |
| Range | 39.00 - 47.00 | 40.00 - 49.00 |  |
| UHDRS TMS score |  |  | 0.887^b^ |
| Mean (SD) | 5.67 (4.74) | 5.79 (3.74) |  |
| Range | 0.00 - 22.00 | 0.00 - 21.00 |  |
| CAP score |  |  | 0.860^b^ |
| Mean (SD) | 85.78 (10.89) | 85.30 (13.01) |  |
| Range | 62.56 - 104.01 | 62.10 - 125.12 |  |
| DBS score |  |  | 0.912^b^ |
| Mean (SD) | 304.20 (59.12) | 302.88 (51.89) |  |
| Range | 179.00 - 401.00 | 185.00 - 443.00 |  |
| HD-ISS staging |  |  | 0.154^a^ |
| 0 | 21 (30.88%) | 5 (16.67% |  |
| 1 | 27 (39.71%) | 13 (43.33%) |  |
| 2 | 18 (26.47%) | 8 (26.67%) |  |
| 3 | 2 (2.94%) | 4 (13.33%) |  |
| **Psychiatric Characteristics** | | | |
| Current depression |  |  | **< 0.001^a^** |
| No | 12 (40.00%) | 68 (100.00%) |  |
| Yes | 18 (60.00%) | 0 (0.00%) |  |
| BDI-II score |  |  | 0.064^b^ |
| Mean (SD) | 9.57 (11.32) | 6.01 (7.19) |  |
| Range | 0.00 - 46.00 | 0.00 - 27.00 |  |
| BDI-II cut-off |  |  | 0.739^a^ |
| Clinically elevated | 8 (26.67%) | 16 (23.53%) |  |
| Not clinically elevated | 22 (73.33%) | 52 (76.47%) |  |
| HADS-D score |  |  | **0.020^b^** |
| Mean (SD) | 4.73 (5.10) | 2.62 (3.55) |  |
| Range | 0.00 - 19.00 | 0.00 - 14.00 |  |
| HADS-D cut-off |  |  | 0.213^a^ |
| Clinically elevated | 7 (23.33%) | 9 (13.24%) |  |
| Not clinically elevated | 23 (76.67%) | 59 (86.76%) |  |
| Current mood medication use |  |  | **0.001^a^** |
| No | 16 (53.33%) | 57 (83.82%) |  |
| Yes | 14 (46.67%) | 11 (16.18%) |  |
| History of suicidal ideation |  |  | **0.041^a^** |
| No | 19 (63.33%) | 56 (82.35%) |  |
| Yes | 11 (36.67%) | 12 (17.65%) |  |
| History of suicide attempts |  |  | 0.288^a^ |
| No | 27 (90.00%) | 65 (95.59%) |  |
| Yes | 3 (10.00%) | 3 (4.41%) |  |

Significant p values (≤ 0.05) are shown in bold. SD = standard deviation. CAG = cytosine-adenine-guanine. CAP = CAG-Age Product. DBS = Disease Burden Score. BDI-II = Beck Depression Inventory, 2nd Edition. HADS-D = Hospital Anxiety and Depression Scale, Depression Subscale. ^a^Pearson’s Chi-squared test ^b^Linear Model ANOVA

##### Depression history

SI.Table 2 Depression history classification details

|  | **History of depression (*n* = 30)** |
| --- | --- |
| **Remitted depression** | 12 (12.2%) |
| 1 remitted episode | 11 (91.67%) |
| 2 remitted episodes | 1 (8.33%) |
| **Current Episode of Depression** | 18 (18.4%) |
| First episode of depression | 17 (17.3%) |
| Recurrent episode of depression | 1 (1.0%) |
| Median duration of current episode - years (IQR), range | 4 (2.3 - 6.7), 0.3 - 37.7 |
| **ICD 10 diagnostic category** |  |
| F32.9 (depressive episode, unspecified) | 29 (96.7%) |
| F32.2 (severe depressive episode without psychotic symptoms) | 1 (3.3%) |

ICD = International Statistical Classification of Diseases and Related Health Problems, 10th Revision

##### Mood medication use

Medications were used commonly for depression, anxiety, post-traumatic symptoms, insomnia, and irritability, with 25 participants were taking mood medications and 23 (92%) taking only one. Consistent with existing research using the same cohort(11,12,61–66), we included a covariate to account for medication use (SI.Table 3).

SI.Table 3 Mood medication use

| **Medication** | ***N*** | ***n* (depression history)** | ***n* (no depression history)** | **Duration mean (sd), range – years** | **Dose mean (sd), range – mgs** | **Indication** | **Regime** |
| --- | --- | --- | --- | --- | --- | --- | --- |
| **SSRI** |  |  |  |  |  |  |  |
| Citalopram | 11 | 5 | 6 | 2.11 (1.69), 0.40-5.73 | 20.9 (8.3), 10.0-40.0 | anxiety (27%), depression (73%) | once daily (100%) |
| Escitalopram | 2 | 1 | 1 | 2.44 (3.46), 0.00-4.89 | 10.0 (0.0), 10.0-10.0 | depression (100%) | once daily (100%) |
| Fluoxetine | 1 | 1 | 0 | 3.87 (0.00), 3.87-3.87 | 20.0 (0.0), 20.0-20.0 | depression (100%) | once daily (100%) |
| **Non-SSRI** |  |  |  |  |  |  |  |
| Amitriptyline | 1 | 1 | 0 | 13.78 (0.00), 13.78-13.78 | 50.0 (0.0), 50.0-50.0 | post-traumatic stress (100%) | once daily (100%) |
| Bupropion | 1 | 0 | 1 | 0.66 (0.00), 0.66-0.66 | 150.0 (0.0), 150.0-150.0 | anxiety (100%) | once daily (100%) |
| Mianserin | 1 | 0 | 1 | 0.41 (0.00), 0.41-0.41 | 30.0 (0.0), 30.0-30.0 | depression (100%) | once daily (100%) |
| Mirtazapine | 1 | 1 | 0 | 1.04 (0.00), 1.04-1.04 | 15.0 (0.0), 15.0-15.0 | depression (100%) | once daily (100%) |
| Venlafaxine | 2 | 2 | 0 | 0.74 (0.33), 0.51-0.98 | 93.8 (26.5), 75.0-112.5 | depression (100%) | twice daily (50%), once daily (50%) |
| **Antipsychotic** |  |  |  |  |  |  |  |
| Aripiprazole | 1 | 1 | 0 | 0.31 (0.00), 0.31-0.31 | 15.0 (0.0), 15.0-15.0 | depression (100%) | once daily (100%) |
| Olanzapine | 1 | 1 | 0 | 2.06 (0.00), 2.06-2.06 | 2.5 (0.0), 2.5-2.5 | irritability (100%) | once daily (100%) |
| **Benzodiazepine** |  |  |  |  |  |  |  |
| Bromazepam | 1 | 0 | 1 | 3.24 (0.00), 3.24-3.24 | 1.5 (0.0), 1.5-1.5 | insomnia (100%) | once daily (100%) |
| Oxazepam | 1 | 0 | 1 | 0.41 (0.00), 0.41-0.41 | 10.0 (0.0), 10.0-10.0 | anxiety (100%) | twice daily (100%) |
| Temazepam | 1 | 1 | 0 | 14.83 (0.00), 14.83-14.83 | 10.0 (0.0), 10.0-10.0 | sleep disorder (100%) | once daily (100%) |

SSRI = Selective Serotonin Reuptake Inhibitors. All doses in mg, unless otherwise stated.

#### Principal component analysis of scanner effects

Given that two different scanner systems were used, framewise displacement output from MRIQC v22.0.6(16) assessed for significant differences in scanner output. Visual inspection (SI.Fig 1) and both Kolmogorov–Smirnov test and t-tests assessed significant group differences.

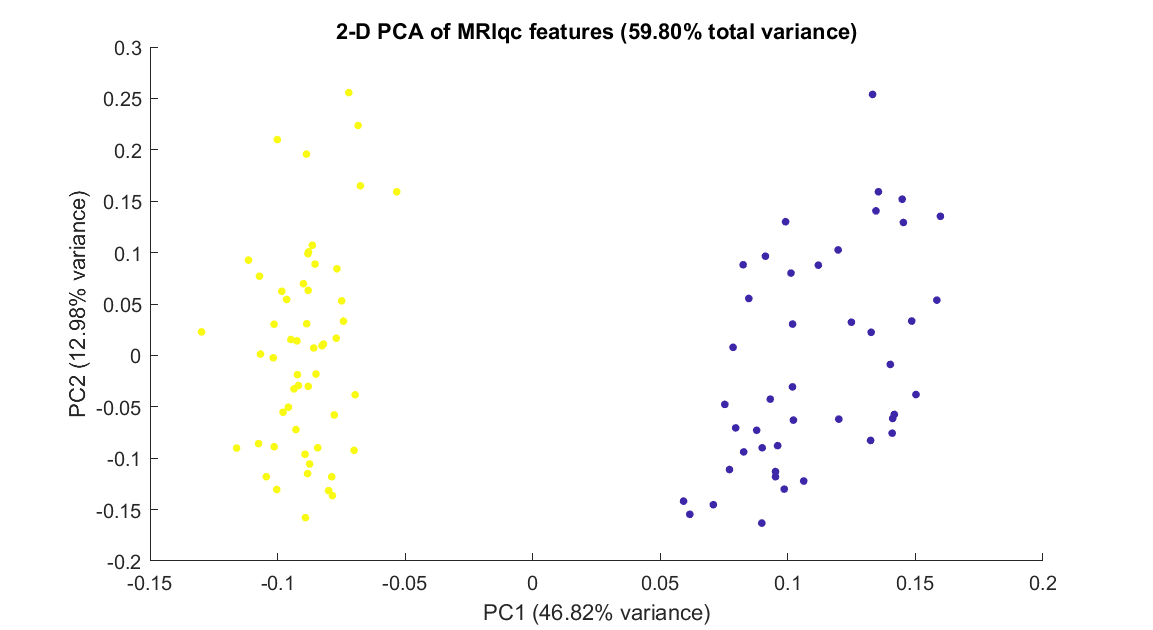

SI.Fig. 1 Principal component analysis of scanner effects | The first two principal components are plotted, with data points colored by scanner type, blue = Siemens; yellow = Philips.

Framewise displacement differed significantly between scanner types, with Philips scanners showing higher mean FD than Siemens scanners (M = 0.224 vs. 0.177 mm; *t*(96) = -2.41, *p* = .018; Kolmogorov-Smirnov *D* = 0.304, *p* = .017). The first principal component, explaining 46.82% of variance in MRIQC quality metrics, was included as a covariate in analyses.

##### Structural connectivity differences

SI.Table 4 DMN and striatum edge strength analyses

| **Connection** | ***t*(96)** | **Pcorr** | **Cohen’s *d*** |
| --- | --- | --- | --- |
| CAU_L_ – CAU_R_ | -0.41 | .87 | -0.13 |
| CAU_L_ – PU_L_ | -0.29 | .87 | -0.09 |
| CAU_L_ – PU_R_ | -1.42 | .74 | -0.44 |
| CAU_L_ – HPC_L_ | 1.83 | .74 | 0.58 |
| CAU_L_ – HPC_R_ | 1.34 | .74 | 0.42 |
| CAU_L_ – PCC | -0.50 | .87 | -0.16 |
| CAU_L_ – MPFC | 0.24 | .87 | 0.08 |
| CAU_R_ – PU_L_ | -0.43 | .87 | -0.13 |
| CAU_R_ – PU_R_ | -0.13 | .90 | -0.04 |
| CAU_R_ – HPC_L_ | 0.95 | .86 | 0.30 |
| CAU_R_ – HPC_R_ | -0.45 | .87 | -0.14 |
| CAU_R_ – PCC | -1.24 | .77 | -0.39 |
| CAU_R_ – MPFC | -1.47 | .74 | -0.46 |
| PU_L_ – PU_R_ | -0.47 | .87 | -0.15 |
| PU_L_ – HPC_L_ | 1.82 | .74 | 0.57 |
| PU_L_ – HPC_R_ | -0.74 | .86 | -0.23 |
| PU_L_ – PCC | -0.28 | .87 | -0.09 |
| PU_L_ – MPFC | 0.75 | .86 | 0.24 |
| PU_R_ – HPC_L_ | -0.24 | .87 | -0.07 |
| PU_R_ – HPC_R_ | 0.98 | .86 | 0.31 |
| PU_R_ – PCC | 0.69 | .86 | 0.22 |
| PU_R_ – MPFC | -0.55 | .87 | -0.17 |
| HPC_L_ – HPC_R_ | 0.78 | .86 | 0.25 |
| HPC_L_ – PCC | -0.20 | .87 | -0.06 |
| HPC_L_ – MPFC | 1.46 | .74 | 0.46 |
| HPC_R_ – PCC | 1.61 | .74 | 0.50 |
| HPC_R_ – MPFC | 0.89 | .86 | 0.28 |
| PCC – MPFC | -0.88 | .86 | -0.28 |

P_corr_ = corrected p-values for multiple comparisons (FDR). No node survived correction. Cohen's d was used as a measure of effect size. - = undirected connection; MPFC = Medial Prefrontal Cortex; PCC = Posterior Cingulate Cortex; HPC_L_ = Left Hippocampus; HPC_R_ = Right Hippocampus; CAU_L_ = Left Caudate; PU_L_ = Left Putamen; PU_R_ = Right Putamen.

SI.Table 5 DMN and striatum node strength analyses

| **Region** | **t(96)** | **P_corr_** | **Cohen’s *d*** |
| --- | --- | --- | --- |
| CAU_L_ | -0.25 | .98 | -0.08 |
| CAU_R_ | -0.44 | .98 | -0.14 |
| PU_L_ | -0.07 | .98 | -0.02 |
| PU_R_ | 0.03 | .98 | 0.01 |
| HPC_L_ | 2.08 | .34 | 0.65 |
| HPC_R_ | 0.96 | .91 | 0.3 |
| PCC | -0.06 | .98 | -0.02 |
| MPFC | -1.28 | .82 | -0.4 |

P_corr_ = corrected p-values for multiple comparisons (FDR). No node survived correction. Cohen's d was used as a measure of effect size. MPFC = Medial Prefrontal Cortex; PCC = Posterior Cingulate Cortex; HPC_L_ = Left Hippocampus; HPC_R_ = Right Hippocampus; CAU_L_ = Left Caudate; PU_L_ = Left Putamen; PU_R_ = Right Putamen.

##### Regional effective connectivity changes

Between region effective connectivity changes between HDGECs with and without a history of depression

| **From → To** | **Valence (No Dep)** | **Valence (Dep)** | **Posterior expectation (difference)** | **95% CI** |
| --- | --- | --- | --- | --- |
| CAU_L_ → PU_L_ | Inhibitory | Inhibitory | 0.102 | [-0.020, 0.225] |
| CAU_L_ → PU_R_ | Excitatory | Inhibitory | -0.088 | [-0.196, 0.021] |
| CAU_R_ → CAU_R_ | Inhibitory | Inhibitory | 0.117 | [0.002, 0.232] |
| CAU_R_ → PU_R_ | Inhibitory | Inhibitory | 0.093 | [-0.021, 0.208] |
| CAU_R_ → HPC_L_ | Inhibitory | Inhibitory | 0.165 | [0.030, 0.300] |
| CAU_R_ → HPC_R_ | Inhibitory | Inhibitory | 0.157 | [0.021, 0.293] |
| CAU_R_ → PCC | Inhibitory | Excitatory | 0.193 | [-0.027, 0.413] |
| CAU_R_ → MPFC | Inhibitory | Inhibitory | 0.410 | [0.226, 0.594] |
| PU_L_ → CAU_L_ | Inhibitory | Excitatory | -0.127 | [-0.257, 0.004] |
| PU_R_ → CAU_L_ | Inhibitory | Inhibitory | -0.121 | [-0.273, 0.030] |
| PU_R_ → CAU_R_ | Inhibitory | Inhibitory | -0.104 | [-0.241, 0.033] |
| PU_R_ → HPC_R_ | Inhibitory | Inhibitory | -0.192 | [-0.328, -0.056] |
| HPC_L_ → HPC_L_ | Inhibitory | Inhibitory | 0.092 | [-0.021, 0.206] |
| HPC_L_ → PCC | Inhibitory | Inhibitory | 0.248 | [0.020, 0.476] |
| HPC_R_ → CAU_L_ | Inhibitory | Inhibitory | -0.088 | [-0.205, 0.028] |
| MPFC → PU_L_ | Excitatory | Inhibitory | -0.065 | [-0.140, 0.009] |

Dep = depression; MPFC = Medial Prefrontal Cortex; PCC = Posterior Cingulate Cortex; HPC_L_ = Left Hippocampus; HPC_R_ = Right Hippocampus; CAU_L_ = Left Caudate; PU_L_ = Left Putamen; PU_R_ = Right Putamen; CI = confidence interval.

##### Leave-one-out cross validation

Leave-one-out cross validation was performed on connections that demonstrated significant associations with depression symptom severity. SI.Fig. 2 shows the individual connections that met significance for leave-one-out cross validation for either HADS-D or BDI-II clinical cut-off for significant depression symptoms.

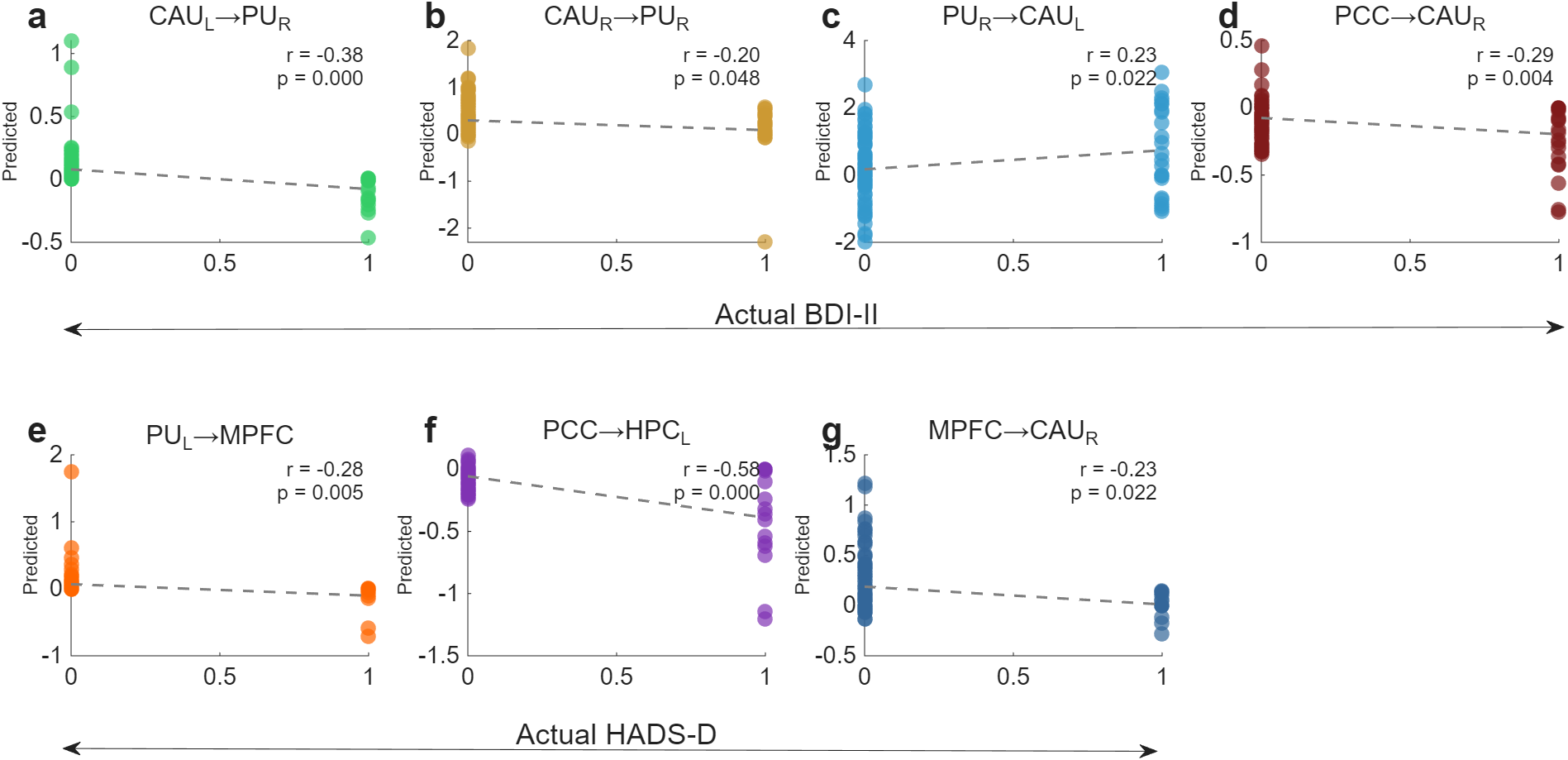

**SI.Fig. 2 Leave-one-out cross-validation correlations between effective connectivity and depression severity for individual connections |** Each panel shows correlations between predicted and actual depression scores on Beck depression inventory, 2nd edition [bdi-ii] **(top; a-d)** or hospital anxiety and depression scale, depression subscale [HADS-D] **(bottom; e-g)**. Correlation coefficients (*r*) and p-values are displayed for each analysis. MPFC = Medial Prefrontal Cortex; PCC = Posterior Cingulate Cortex; HPC_L_ = Left Hippocampus; HPC_R_ = Right Hippocampus; CAU_L_ = Left Caudate; PU_L_ = Left Putamen; PU_R_ = Right Putamen.
